# Phasor-Based Multi-Harmonic Unmixing for In-Vivo Hyperspectral Imaging

**DOI:** 10.1101/2022.03.31.486485

**Authors:** Alexander Vallmitjana, Paola Lepanto, Florencia Irigoin, Leonel Malacrida

## Abstract

Hyperspectral imaging (HSI) is a paramount technique in biomedical science, however, unmixing and quantification of each spectral component is a challenging task. Traditional unmixing relies on algorithms that need spectroscopic parameters from the fluorescent species in the sample. The phasor-based multi-harmonic unmixing method requires only the empirical measurement of the pure species to compute the pixel-wise photon fraction of every spectral component. Using simulations, we demonstrate the feasibility of the approach for up to 5 components and explore the use of adding a 6th unknown component representing autofluorescence. The simulations show that the method can be successfully used in typical confocal imaging experiments (with pixel photon counts between 10^1^ and 10^3^). As a proof of concept, we tested the method in living cells, using 5 common commercial dyes for organelle labeling and we easily and accurately separate them. Finally, we challenged the method by introducing a solvatochromic probe, 6-Dodecanoyl-N,N-dimethyl-2-naphthylamine (LAURDAN), intended to measure membrane dynamics on specific subcellular membrane-bound organelles by taking advantage of the linear combination between the organelle probes and LAURDAN. We succeeded in monitoring the membrane order in the Golgi apparatus, Mitochondria, and plasma membrane in the same in-vivo cell and quantitatively comparing them. The phasor-based multi-harmonic unmixing method can help expand the outreach of HSI and democratize its use by the community for it does not require specialized knowledge.

## 1. Introduction

Nowadays every commercial microscope offers Hyperspectral imaging (HSI) in different acquisition configurations [1,2]. A significant advantage of HSI over traditional epifluorescence detection with bandpass filters is that it inherently solves the problem of bleed-through between channels by collecting the entire spectrum. In HSI all photons are collected within a spectral range increasing the total signal to noise ratio. Most commercial instruments cover the spectral range from 400 to 700 nm, making HSI ideal for *in vivo* imaging in the visible range without adding constraints by selecting a spectral band. Traditionally, microscope manufacturers offer different unmixing approaches, such as classical linear unmixing for supervised unmixing. These unmixing methods rely on *a priori* knowledge of the fluorophores involved in the mixture; to model their fluorescence emission, they require knowledge of the fluorescence emission parameters, such as its maxima and full-width-at-half-maximum. Under circumstances where this information is not available, supervised methods cannot proceed with the unmixing, an example of this being the solvatochromic probe LAURDAN [3], a constraint that hinders the use and adoption of HSI. Some of the methods used for unsupervised unmixing are principal component analysis, independent component analysis or linear discriminant analysis [4–6], which although having the advantage of being unsupervised, are highly technical and harder to implement in a comprehensive manner for the end user. Thus, a simple and model-free method is needed for these circumstances where traditional unmixing cannot be applied due to complex photophysics, molecular interactions, or autofluorescence [3,7–12]. The method we propose here is very close to a supervised linear unmixing, but it operates in the phasor space. A comparison of our method to linear unmixing is shown in the results section.

The spectral phasor approach is a model-free method initially used for unmixing three components [7]. It is a powerful tool for visualizing and analyzing multiple-component spectral images. It has its roots in the study of alternating current systems from which it was borrowed to represent fluorescence lifetime measurements [13–15]. Its extension from the lifetime domain to the spectral domain is relatively recent, with the first description in 2012 [7]. Since then, it has seen many applications [3,12,16] and has allowed the development of important hardware implementations [17–19] and post-processing techniques [20,21].

The spectral phasor transform is an operation that computes two quantities from a given photon distribution as a function of wavelength (e.g. an emission spectrum). These two quantities capture the shape and position of the distribution within the spectral window it was acquired in. Since Weber’s initial work in 1981 [22], these two variables are named *S* and *G.* They can be seen as a pair of coordinates (or a complex number) in the newly transformed phasor space. Therefore, every possible photon distribution has its phasor coordinates and hence a representative location in the phasor space. When performing spectral imaging, each pixel of the image will have its spectral photon distribution, and therefore each pixel will have its own phasor coordinates, mapping to the phasor space. The phasor plot is then a 2D histogram of the S and G coordinates for all pixels in the image, ranging the wavelength period (λ_0_ to λ_f_) in a 2π phasor plot with a radius of 1 (Figure 1B).

**Figure 1.**
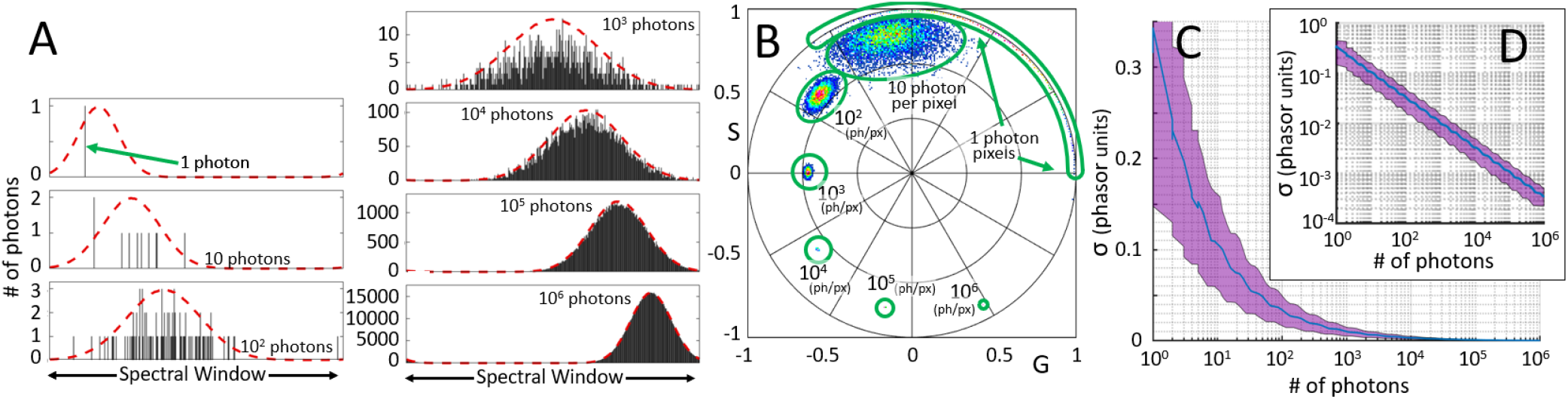
The Spectral Phasor Transform. A) Example of seven simulated photon histograms for an increasing number of photons drawn from the underlying normal distribution depicted in dotted lines. For exemplification, the mean position of the distribution increases monotonically, and the variance grows in the first three cases and decreases in the last three. B) For each of the seven underlying distributions, a 100×100 image is simulated (10^4^ pix) by drawing from the distributions and mapping the resulting histograms to the phasor plot. C) The average standard deviation of each phasor cloud is plotted against the number of photons (10^2^ phasor clouds with 10^4^photon histograms each). D) Log-log plot of the previous representation.

Viewing the spectral phasor plot as a polar plot, the angular position captures the mean position of the photon distribution in the spectral window. More precisely, the phase angle is linear with the mean wavelength (the center of mass of the distribution). At the same time, the distance to the origin on the phasor plot captures the broadness of the photon distribution. The modulation measures the width of the distribution: an infinitely narrow photon distribution lies precisely on the unity circle, whereas a flat distribution lies at the origin of coordinates. As an example, **Figure** 1A shows seven examples of photon sampling from distributions with varying mean wavelength and spectral broadness. **Figure** 1B shows the phasor plots of 10^4^ pixels using the previous underlying distributions at an increasing number of photons sampled per pixel.

Given an underlying photon distribution, the number of photons collected determines how closely the obtained spectra resemble the underlying photon distribution. In other words, the higher the number of photons, the closer the phasor position is to the *true* phasor position of the underlying photon distribution. An in-depth study on phasor scattering statistics and the different factors involved in the error can be found in the work by Cutrale et al. [16]. The result is that the variance of the data clouds on the phasor space is inversely proportional to the square root of the number of photons (**Figure** 1C and D).

At the core of the phasor transforms are the sine and cosine functions that act as filters to the spectral distribution that is being measured. The transform consists in fitting these two trigonometric functions in the spectral window and modulating the spectral distributions to capture the shape and position of the spectral distribution. It is analogous to computing the 1^st^ even and odd coefficients of the Fourier series of this photon distribution. Nevertheless, one can fit the trigonometric functions twice, thrice, or many times in the spectral window, capturing more complex distributions such as bi-modal or tri-modal. Transforming in this way gives rise to new coefficients of the Fourier series, and the values obtained are named the higher harmonics of the phasor transform.

In an experimental measurement, a sample containing several fluorescent species produces a particular photon distribution and establishing the relative contributions of these species to the distribution is of particular interest. Because of the two-dimensionality of the phasor space, only up to three species can be resolved using just the phasor coordinates of the known species [7]. If one computes higher harmonics of the phasor transform, this introduces new relationships to the system, which allows resolution of more than three components. An example of this operation can be seen in **Figure** 2, where we construct 5 pure spectral components, and simulate two mixtures of them such that the 1st harmonic maps the two mixtures to the same coordinates on the phasor plot (**Figure** 2B). The second harmonic phasor transform of the same two mixtures maps to different locations, therefore, resolving the ambiguity (**Figure** 2C).

**Figure 2.**
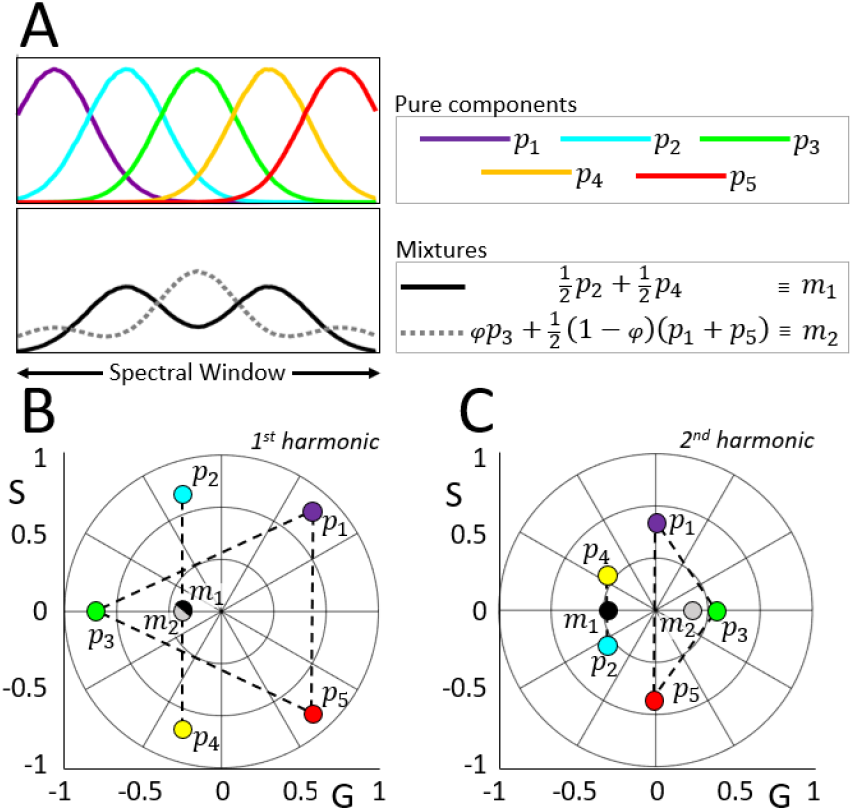
Phasor ambiguity unmixed by 2^nd^ harmonic. A) Five spectral photon distributions are used to compose two fractional mixtures. B) The two mixtures are chosen such that they fall on the exact same spot on the phasor plot. C) The 2^nd^ harmonic unmixes the positions, allowing computation of the relative fractions using the combined relationships of the two harmonics. For this particular example, due to the geometry of the pentagon, the fractions of *m_2_* happen to contain the irrational number *φ*, the golden ratio.

In this paper, we show how we can resolve up to 5 individual spectral components using two harmonics of the phasor transform with a phasor-based linear unmixing method. This method is an extension of our previously developed unmixing technique used in lifetime phasor data [23] and is here extended to higher harmonics and a higher number of components. We use simulations to establish a relationship between the number of photons required and the accuracy in the recovery of the fractional components. In addition, we show how the more realistic situation, in which the sample contains additional unknown fluorescent components, introduces a minimum error in the accuracy. The spectral phasor representation and unmixing has high potential in solving spectroscopic problems such as the ones that arise with solvatochromic dyes. For this reason, we challenged our method to understand the membrane’s physical order and dynamics within living cells by combining LAURDAN fluorescence with organelle-specific target fluorescent labels. This is a difficult and valuable question due to the impossibility to inform specific organelle membrane dynamics with the current methods [24]. Applying the multi-harmonic phasor unmixing, we were capable of obtaining the fraction of each labeled organelle and the membrane fluidity in a system of 5 spectral components.

LAURDAN is a membrane probe sensitive to its dipolar environment and can relax dipolar molecules around it [10]. In membranes, LAURDAN is used to measure supramolecular membrane organization, often known as fluidity or lipid order [25]. Interestingly, its fluorescence changes (either spectra and lifetime) involve the physical order of the membrane due to its thermodynamic status and the effects of internal and/or external components, such as cholesterol, proteins, ionic strength, pH, water activity, and metabolites/drugs interacting with the membranes [10,26–29]. All these variables modify the membrane physical parameters differentially, making it impossible to predict the actual environment for LAURDAN in systems such as *in vivo* cells. Thus, a method based on the phasor approach analysis of the spectral data without proposing a model such as in the generalized polarization function or any other unmixing, is of great value [3].

## 2. Results

### 2.1 Simulations

#### 2.1.1 Unmixing error as a function of photon counts

We ran a set of simulations to establish the performance of the method as a function of the number of photons acquired in an experimental measurement. We downloaded the spectral emission profile of a set of 5 widely used fluorophores from the Chroma spectra-viewer online utility (Figure 3A) and we computed the spectral phasor positions of these components for the first 3 harmonics using a spectral window of 450nm to 700nm (first harmonic shown in Figure 3B).

**Figure 3.**
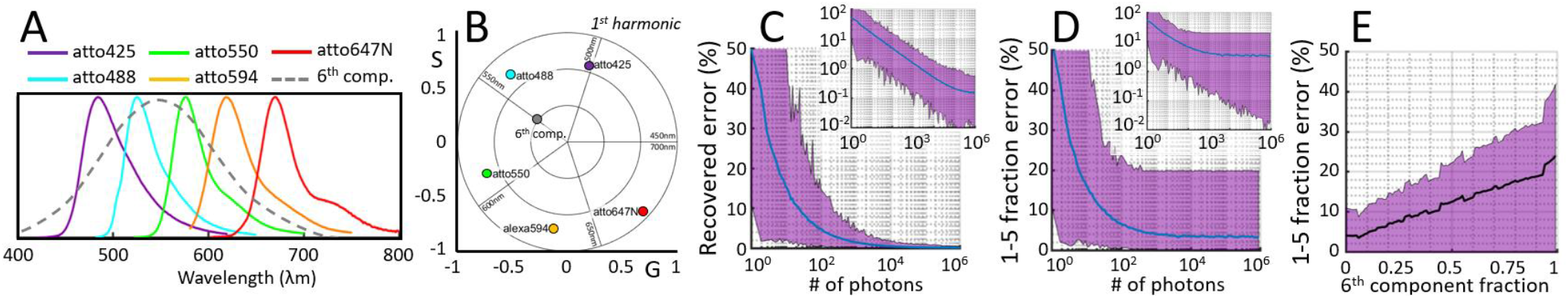
Simulation results. A) Emission spectra of five known fluorescent probes plus a constructed 6^th^ component (used in the second simulation). B) The phasor positions of the depicted emission spectra. C) A total of 400,000 simulations were carried out. In each case, random fractions and random numbers of photons for the 5 components were used. After recovery of the fraction of each component, the mean error between the recovered and the real fraction is depicted as a function of the number of photons. Inset depicts the same data on a log-log scale. D) A new set of simulations was carried out in which a 6^th^ component is added to the generated random mixtures and again the mean error of the recovered fractions is plotted against the number of photons. E) In the same simulation, the mean error is plotted as a function of the fraction of the 6^th^ component. In all cases, shaded areas correspond to the maximum and minimum error intervals.

A total of 4×10^5^ spectral measurements were simulated, with the spectra containing from 1 photon to 10^6^ photons. For each simulated experiment, we randomly generated a measured spectrum using Monte-Carlo sampling with a randomly generated set of fractions. This approach provided a ground truth of the fractions together with a spectral measurement which was then unmixed using our phasor-based multi-harmonic unmixing method. For each simulated experiment, we then computed the mean, maximum, and minimum between the absolute value of the difference between the 5 recovered fractions and the 5 known true fractions (Figure 3C).

This simulation reveals a linear relationship between the order of magnitude of the number of photons and the order of magnitude of the error committed in the recovered fractions. The error is considerably higher when dealing with a low number of counts: 15% error in the recovered fractions when operating in the range of 10 photons per pixel, which decreases to 5% with 100 photons per pixel and 1% at 10^3^ photons per pixel (Figure 3C).

#### 2.1.2 Unmixing error in presence of autofluorescence

The second set of simulations were designed to study the effect of unknown components that may be present in a sample. This case is much closer to a real experiment in which, in addition to the known dyes, there is a contribution from some naturally fluorescing molecules contributing to a background component, for example autofluorescence in cells. Because it is a combination of many different components, autofluorescence tends to show up as a broad distribution, i.e. low modulation, centered around the middle of the visible spectrum (see Supplementary Figure 1), in relation to our simulated spectral phasor plot (ranging from 450 nm to 700 nm), in the vicinity of the transition from the 2nd to 3rd quadrant. Thus, we modeled autofluorescence with a gaussian function, centered at 550 nm, with a large broadness: 3 standard deviations of 180 nm (dashed line in Figure 3A, gray dot in Figure 3B).

The simulation was performed similarly to the previous case but with the added 6th component representing autofluorescence, when generating the data, but not when recovering the fractions. We randomly generated spectra using linear combinations of the 6 fractions (5 known components plus the 6th representing autofluorescence) and we then ran the phasor-based linear unmixing algorithm to attempt to recover the fraction of the known 5 components without considering the presence of the 6th.

As one would expect, this process introduces an error to the recovered fraction, but interestingly it establishes a minimum accuracy that cannot be improved by having brighter samples (figure 3D). Compared to the values exemplified in the previous section, with 10 photons, the recovered error is the same as before at 15%. This error is simply due to a lack of photons, but now with 10^3^ photons, the error is 4% instead of 1%, manifesting the effect of the hidden component. This error now remains practically constant at 3% with 10^6^ photons. Furthermore, this error is dependent on the relative proportion of photons originating at the 6th unknown component; if the signal from the known components is very low compared to the 6th, the error is greatly increased (figure 3E).

#### 2.1.3 Comparison to linear unmixing

Using the same set of 5 spectra (Figure 3A), we generated a total of 10M simulated mixtures drawing uniform random fractions of each component and random number of photons out of a logarithmic distribution. We solved each of the simulated 28ch-sampled spectra with our multi-harmonic phasor method and solved also using linear unmixing (both using the same least squares algorithm to solve the system of linear equations). The results showed that with high enough number of photons (above 10^3^) both methods produced the same results (fraction errors below 10^-4^). In regimes of lower photon counts, classical linear unmixing performed better (up to 0.03 fraction difference) when recovering low fractional values (under 0.66) and multi-harmonic phasor unmixing performed better (up to 0.01 fraction difference) when recovering higher fractional values (above 0.66). These results suggest that the best approach should be an ensemble method of the two where each is favored depending on the recovered fractional values. See Supplementary Figure 3 for the full results of the comparison.

### 2.2 Organelle-specific stained cells

We tested our phasor-based multi-harmonic unmixing method on living cells loaded with several organelle-specific markers. The first proof-of-concept experiment used 5 of the most common used commercial organelle markers (Figure 4) for the nucleus (Comp.1), lysosomes (Comp.2), Golgi apparatus (Comp.3), mitochondria (Comp.4), and plasma membrane (Comp.5). Cells were first stained with each of the individual components to obtain the phasor positions of the components in each harmonic (see Supplementary Figures 1 and 2). A measurement was then taken on cells loaded with all 5 components simultaneously. In this experiment, we are in a high photon count regime, accumulating all channels peak counts were above 10^3^ (Figure 4A). However, in individual channels we are dealing with regimes in the order of 10^2^ at the most.

**Figure 4.**
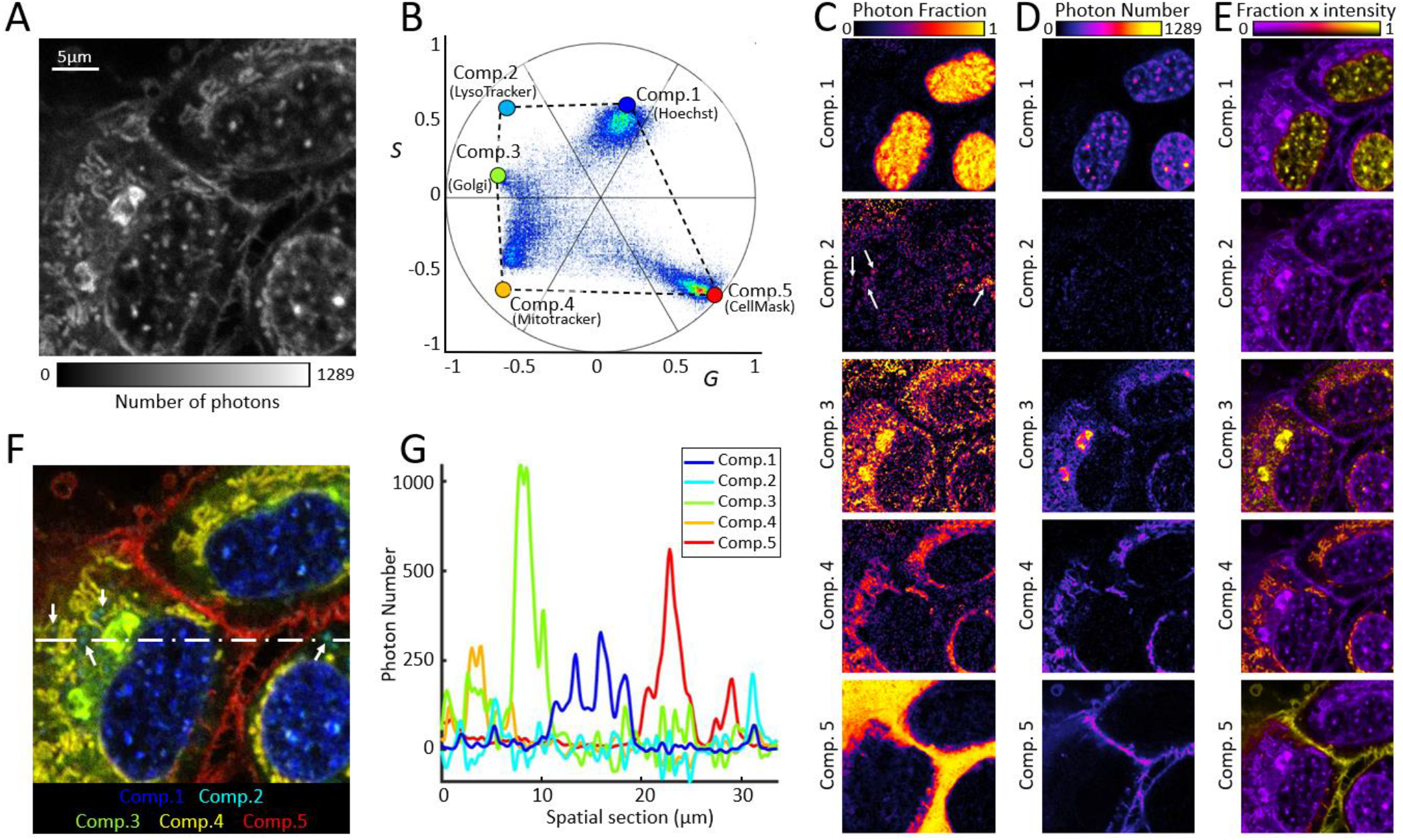
In-vivo results with commercial dyes. A) Intensity image of an example cell stained with Hoechst, LysoTracker Green, CytoPainter Golgi, MitoTracker Orange, and CellMask. B) Spectral phasor plot of the image overlapped with phasor positions of the pure components (previously measured *in vivo*). C) Pixel component fraction of every component unmixed from the image. D) Fractional number of photons unmixed from the image. E) Fraction of photons normalized to the intensity image (A). The lysosome signature, being very dim is hardly observable, for this reason we have included arrows in panel C, component 2, showing examples of its presence, which in turn is depicted in panel F in cyan color. F) Pseudocolor image generated using the unmixing results, with the arrows pointing example lysosomes. The dotted line depicts the line from which the linear intensity profile is extracted (G) to reveal spatial regions where each component dominates.

The phasor-based multi-harmonic unmixing method computed the relative fraction for each component at each pixel which allowed us to construct images with the recovered fraction at each pixel. The resulting images have a high correlation with the expected spatial distribution of the cellular organelles (Figure 4C). We show three possible representations. In the first, for each component, an image is shown where the pixels color-code the recovered photon fraction over unity (Figure 4C). In the second, the recovered pixel photon fraction is multiplied by the photon counts in each pixel, giving an unmixed image with the number of photons assigned to each component (Figure 4D). A final composite uses the intensity image, color-coding by the pixel photon fraction (Figure 4E). Notice that the method performed very well even with the situation of including a fluorophore with a considerably broad emission such as Hoechst, which is technically unmixable with LysoTracker Green when co-excitation with 405/488 nm is used. The final composition of the unmixed components can be generated using pseudo-coloring to depict each component (Figure 4F). It is worth mentioning that because lysosomes are small structures in comparison to some of the other organelles, the phasor plot (Figure 4B), being a pixel-based representation, hides their presence in the bulk of the other pixels. This does not mean that their fraction was incorrectly computed, rather the method is independently applied pixel per pixel and, in whichever pixels that the lysosomes were present, the method still recovered a dominating component assigned to the LysoTracker (arrows, Figure 4F). Because the method interrogates each pixel for the photon fraction of each of the known components, the results do not incorporate nor depend on the relative brightness of each probe, nor the excitation efficiency given the relative powers of the excitation sources.

In a second set of experiments, we included LAURDAN in the staining mixture, which by itself introduced two components due to its spectral shift as a function of local molecular conditions. The final goal of this experiment was to recover the membrane fluidity (or membrane order) indicated by the LAURDAN fluorescence in each of the subcellular organelles included in the study. As before, we first measured experimentally the phasor positions of the individual dyes in separate in-vivo experiments (see Supplementary Material), among which are the two extreme locations of the whole range of emission of LAURDAN in membrane models (Figure 5B). Even in this case, the images had relatively few counts (maximum photon value of 671, fig 5A), we successfully unmixed the 5 components fig 5C-F.

**Figure 5:**
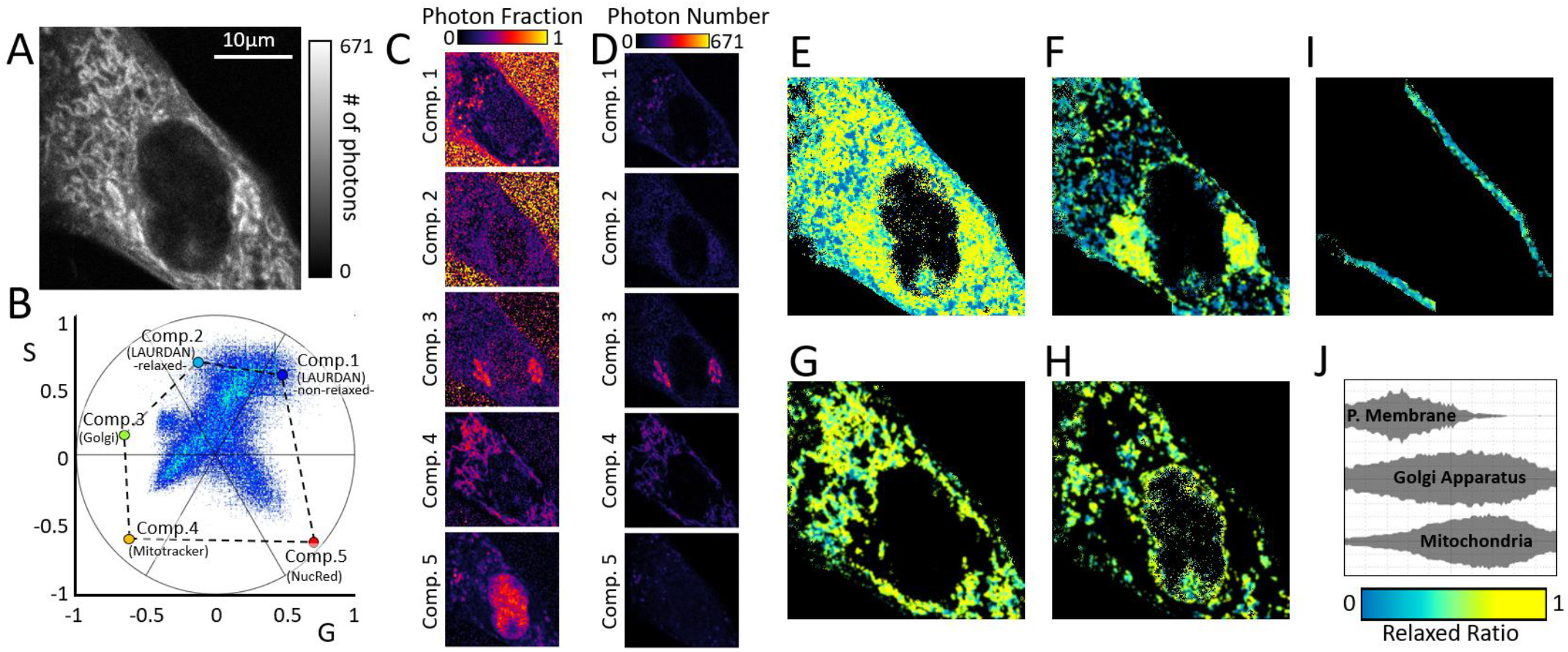
*In vivo* results. A) Intensity image of an example cell stained with LAURDAN, CytoPainter Golgi, MitoTracker Orange, and NucRed. B) Spectral phasor of the image overlapped with phasor positions of the pure components (previously measured *in-vivo*). C) Fraction of photons of each component unmixed from the image. D) Number of photons unmixed from the image. E) LAURDAN relaxed ratio unmixed from the image (intensity threshold set at 100 counts). F-I) LAURDAN relaxed ratio on pixels dominated by each of the remaining components: F) Component 3 (Golgi), G) Component 4 (Mitochondria), H) Component 5 (Nuc Red), and I) Plasma membrane (segmented by erosion of (E)). J) Violin plots of the relaxed ratio values from the pixels segmented in the image.

The power and limitations of the method can be observed in Figure 5. For example, in situations of extremely low counts, in the order of 1 to 10 photons (e.g. outside the cell in Figure 5C), the system simply cannot deal with such sparse spectral intensity profiles, the phasor locations have a huge variance (exemplified in Figure 1B) and therefore the error is also large. As an example, the areas outside the cell are showing extremely high variance in the resulting fractions, many times producing impossible combinations of fractions (such as above unity or below zero). Nevertheless, at a photon regime of a few dozen counts (Figure 5C, inside the nucleus), the method is already picking up the significant presence of the 6th component. It is worth mentioning that if one color-codes the photon fraction (Figure 5D) instead of the pure fraction (Figure 5C), the presence of low-photon count components is masked by the higher counts of the other regions of the image (the nucleus is not visible in Figure 5D, last panel).

To study the physical state of the membrane by the LAURDAN fluorescence, we defined the LAURDAN relaxed ratio as the pixel-wise ratio between the relaxed LAURDAN fraction over the sum of the fractions of the two LAURDAN components. It is equivalent to removing the pixel-wise contribution of all other components and leaving only the LAURDAN components, to then renormalize to the sum of the fractions of LAURDAN’s two components (Figure 5E). A similar exercise can then be done for the other components: we exclude the fraction of LAURDAN, and then renormalize the remaining fractions to their sum. We then used this renormalized fractional contribution to segment the individual organelle pixels, by assigning at each pixel the component with the highest fraction. This calculation allowed us to go back and analyze the LAURDAN relaxed ratio in the pixels belonging to each individual organelle, represented in fig 5F. The results indicate that the plasma membrane had a less relaxed (more ordered) profile than the membrane of the mitochondria or Golgi apparatus, suggesting a more rigid membrane. Thus, we were able to determine and compare the relative physical state of different membranes within the same cell.

## 3. Discussion

The phasor-based multi-harmonic unmixing method is proposed to support fluorescence microscopy imaging development, one of the most important technologies in current life science research. However, one of the challenges that remains to expand fluorescence microscopy is the number of simultaneous fluorophores that can be used and the handling of spectral bleed-through in regular laser scanning confocal microscopy. In the extreme case, fluorophores with the same excitation wavelengths and similar emission spectra cannot be used together. A second level of complexity is associated with the use of fluorophores with responses that are sensitive to the environment, having a continuum of states and emission properties [3]. It is for that reason that model-free methods are needed. The spectral phasor approach is a model-free method that allows to unmix up to 3 components on hyperspectral imaging data [7]. Using linear algebra, it is possible to obtain the fraction of components per pixel and use the reciprocity principle to color back the location in the original image considering this data. The use of higher harmonics for many components analysis in time-resolved data was introduced previously by our group [23]. To overcome the limitation of 3 fluorophores as the limit for unmixing using the spectral phasor approach, we implemented the phasor-based multi-harmonic unmixing method to handle up to 2N+1 components with N being the number of harmonics used. This number is dependent on the spectral resolution of the system and, through Nyquist sampling theorem, it corresponds to half the number of channels used in the spectral dimension.

As the simulation in Figure 3C showed, it is possible to recover 5 components using the first and second harmonic equations. For larger numbers of components, there is a need for higher harmonics. An important point to highlight in this discussion is the relevance of the number of photons to the final number of components that can be unmixed. The variance in the phasor coordinates due to lack of photons is enhanced in the higher harmonics, thus the higher the number of photons collected the larger the number harmonics can be accessed and therefore the number of components that can be unmixed [23].

The computational cost of the method is relatively low due to the simple operations involved. At each pixel a system of equations is solved to obtain the fractions. In our MATLAB implementation, solving for 5 components with 2 harmonics, the average computing time is 3.27s/Mpix. As an example, for the 256×256 images used in our figures, the computation of the fractions was less than half a second. For comparison purposes, classical linear unmixing is in the same order of magnitude for it requires similar operations. Phasor unmixing slightly outperforms due to the fact that, the system of equations is only overdetermined at most by one equation, whereas in linear unmixing the system of equations tends to be overdetermined by more equations due to the higher number of channels. For our example case of 5 components, phasor unmixing solves 5 equations, whereas linear unmixing solves for 29 equations (spectra acquired in 28 channels). The same MATLAB implementation of least squares for linear unmixing yields a computational time of 3.47s/Mpix.

Another relevant aspect in the component resolution is the limitation in the phasor positions of the components, e.g. if three fluorophores are in a line. In this case, a measurement in the middle position can be obtained from the emission of the fluorophore the signature of which is in that location, but also as a linear combination of the two fluorophores the signature of which are the extremes of the line. When this happens, the phasor-based multi-harmonic unmixing method cannot properly assign the fractions of the components because the system of equations is undetermined. However, if a fourth component takes place and it is in interaction with the other three occurs at different proportions, this situation helps to resolve the 3 components better because they are pulled at different proportions, i.e. separated in the phasor plot. This is the case for LysoTracker, CytoPainter - Golgi Staining, and Mitotracker Orange (Figure 4). The components are more or less in a line, however, when CellMask Deep Red is in the game, the fourth components can be resolved without confusion.

The proof-of-concept was performed with a selection of the different fluorophores often used as labels for organelles such as the nucleus, Golgi apparatus, lysosomes, mitochondria, and plasma membrane (Supp. Figure 2). Looking at the spectrum of the probes one may notice the heavy overlap between Hoechst and LysoTracker Green, with the DNA probe bleeding throughout the other channels. However, in the spectral phasor plot, each fluorophore takes a position with a particular phase and modulation (Supp. Figure 1). This fingerprint is characteristic in each harmonic and the coexistence of more than one component in the same pixel maps the pixel to a linear combination between the many species that were involved. The spectral phasor data (Figure 4B) showed that the majority of pixels have different ratios of each component as opposed to a single component. The algorithm correctly separated the photons into one of the components and it is clear that component 1 localizes in the nucleus, component 2 in the lysosomes, component 3 in the Golgi apparatus, component 4 in the mitochondria, and component 5 in the plasma membrane. By calculating the fraction times intensity, we can assess the specificity of each probe. One may notice that there are pixels with high component value outside the organelle’s expected location. This is due to the unspecific labeling of other subcellular structures, which could possibly optimized with changes in the labeling concentration and time. However, through the representation of the relative amount of photons of the specific component, it is clear the organelle identification. The final goal of an unmixing algorithm is to produce an image where each of the components can be blended. Figure 4F presented the final outcome of our phasor-based multiharmonic unmixing method, where it is possible to observe the excellent performance of the new approach for unmixing organelle-specific probes.

In our simulation (Figure 3) we assess the role of autofluorescence in underestimating each component. The first point we claim is that autofluorescence can be measured, and if so, it can be roughly included in the analysis system as another component. By doing this, the new component decreases the uncertainty and contributes to every pixel. A second concern is that, realistically, autofluorescence cannot be considered constant across the whole cell or tissue under imaging. This point is important and difficult to overcome. Still, based on the phasor distribution of autofluorescence in our live cells, the variance appears close to gaussian, thus, homogeneous in its component composition. In the case of autofluorescence with a linear trajectory (more than one component with a variable fraction of each one), some other considerations would be needed to model it. Nevertheless, in our experiments with live cells, the role of autofluorescence was relatively small. The peak number of counts using the same laser power in unlabeled cells was on average 83 counts, dismissing the relevance of autofluorescence for our labeling assignment whereas in the labeled cells, our peak counts were in the order of 10^3^.

The final problem to challenge our method is very demanding and not addressable by classical unmixing. We wanted to use a solvatochromic probe to evaluate the membrane’s physical order or dynamics in combination with dyes for specific organelles. LAURDAN is perhaps the probe more studied on *in vitro* and *in vivo* systems [10,30]. Its unique photophysics and lack of affinity for any particular subcellular organelles enable LAURDAN to study the membrane fluidity in the cell as a whole. LAURDAN is located in the same place at the membrane interphase and does not have preferential affinity for one or another phase separation [8]. This fact is key to comparing the different subcellular membranes without the need of utilizing membrane dyes with modification to target a specific organelle. This kind of approach can modify the location of the probe in the membrane, disabling the proper comparison between the results at the different subcellular locations. We aimed to trace the membrane order at different subcellular organelles taking advantage of the linear combination rules of the phasor approach. In previous work, we used LAURDAN and one extra component, such as mCherry labeling a target of interest [31]. But using the phasor-based multi-harmonic unmixing method we were able to label several membranous organelles, with a total of 5 components at the same time (two from LAURDAN and 3 from subcellular organelles), and still recover membrane organization due to LAURDAN photophysics. In Figure 5, the results indicate that component 3 and 4 are associated with Golgi and Mitochondria, respectively, while component 5 is to the nucleus. After assignment to the organelle-specific probe, we performed an analysis of the LAURDAN relaxed ratio as a proxy of the physical order of the membrane. Looking at the whole cell, one may notice that there are a high number of bluish pixels at the perimeter of the cell, which is in line with the expected location for a more ordered plasma membrane. By isolating the plasma membrane region of interest, we quantified the pixel distribution across the relaxed ratio dimension, showing that it is the most ordered membrane in the cell as published before [26]. When comparing the relaxed ratio in the Golgi apparatus to the mitochondria, we observe that the Golgi apparatus distribution is broader and has a slightly lower median, it is possible that this is due to the background contribution of a coarse segmentation. Still it is clear that the Golgi apparatus and the mitochondria are less ordered than the plasma membrane. For component 5, there are two things to be discussed. First, LAURDAN does not fluoresce within the nucleus due to the strong hydrophobic tension, and we see a halo around the nucleus corresponding to the nuclear envelope. The other puncta in the cytoplasm is difficult to assign but certainly can be vesicles where the NucRed is entrapped. In previous studies with LAURDAN on live cells, a significant limitation in our subcellular analysis was the impossibility of discriminating the contribution from the various organelles and studying the membrane order at a specific target organelle. With the three-component analysis using the first harmonic, it was possible to address one at a time, but then the comparison between different sets of LAURDAN plus the third component was impossible. To the best of our knowledge, this is the first report for a method that can address membrane dynamics by LAURDAN fluorescence for several subcellular organelles on *in vivo* cells. This approach opens the possibility to follow *in vivo* membrane organelle interaction and crosstalk during the cell cycle or under stress.

Finally, a relevant aspect to be discussed is the intuitive feature intended for non-expert users. Our method should help in democratizing the use of hyperspectral imaging across life sciences. Other approaches for unmixing require complex models and a certain level of expertise in conjunction with spectral imaging adoption. The only requirement for using the phasor-based multi-harmonic unmixing is to acquire the single components to define the locations of the several phasor components in the different harmonics.

## 4. Methods

### 4.1 Cell Culturing

NIH-3T3 cells (ATCC, CRL-1658) were maintained in DMEM-GlutaMAX (Thermo, #10566016) supplemented with Donor Bovine Serum 10% (Thermo, #16030-074), HEPES (10mM, Thermo, #15630080) and penicillins-treptomycin (100 U/mL, Thermo, #15140122) under a 5% CO2 atmosphere, at 37°C and subcultured using Trypsin-EDTA 0.05% (Thermo, #25300054). For imaging, cells were seeded onto 35mm glass-bottom Petri dishes (Greiner Bio-One, #627870) at a density of 10^5^ cells per compartment one day before staining and imaging.

### 4.2 Cell Staining

The dyes used in this study and the working dilutions were: 6-Dodecanoyl-N,N-dimethyl-2-naphthylamine (LAURDAN, 4.5 μM, Sigma-Aldrich, Merck, #40227), Hoechst 33342 (80 uM, Abcam, ab139483), LysoTracker™ Green DND-26 (50 nM, ThermoFisher, L7526), CytoPainter - Golgi Staining Kit - Green Fluorescence (0.8 nM, Abcam, ab139483), MitoTracker^™^ Orange CMTMRos (0.05 nM, ThermoFisher, #M7510), NucRed^™^ Live 647 ReadyProbes^™^ Reagent (1/20, ThermoFisher, #R37106) and HCS CellMask™ Deep Red Stain (2.5 ug/ml, ThermoFisher, #H32721). Staining with the Golgi dye from the Golgi Staining Kit was performed following the instructions of the provider; briefly, cells were washed in Assay Solution, incubated with Golgi dye for 30 min at 4°C, washed three times in ice-cold complete medium, incubated for 30 min at 37°C in complete medium. Staining of cells with LAURDAN, MitoTracker, ER-Tracker, and NucRed was performed with the concentration indicated by vendor for each dye diluted in complete medium and an incubation of 30 min at 37°C before imaging. The medium with the dyes was maintained throughout the microscopy acquisition.

### 4.3 Multilamellar Vesicle (MLV) generation and staining with LAURDAN

To determine the extremes of LAURDAN trajectory MLV with fluid or liquid order phases were used. The lipids used were the following: 1,2-dioleoyl-sn-glycero-3-phosphocholine (DOPC, Avanti, #850375C), 1,2-dipalmitoyl-sn-glycero-3-phosphocholine (DPPC, Avanti, #850355) and cholesterol (Avanti, #700000). Stock solutions of each lipid were generated in chloroform and used to form a 0.5 mM MLV dispersion of DOPC or a 0.25 mM DPPC plus 0.25 mM cholesterol. The final concentration for LAURDAN in the MLVs dispersion was 0.5 μM. The mixtures were dried with N2 gas followed by vacuum for 30 min. Then, PBS 1X pre-heated to 50°C was added, and the mixtures were subjected to vortexing followed by heating at 50°C for three cycles. An aliquot of each mixture was deposited over a glass-bottom Petri dish and left undisturbed one hour before imaging. Images were acquired at 21°C for the mixture of DPPC-cholesterol and DOPC.

### 4.4 Hyperspectral microscopy Imaging

Hyperspectral acquisitions were performed in a Zeiss LSM 880 microscope, using a Plan-Apochromat 63x/1.4 Oil DIC M27 objective. For excitation, we used four independent single-photon laser lines; a 488 nm Argon laser, together with three diode lasers: 405, 561, and 633 nm. On the emission side, we used a 405 nm long pass dichroic mirror for UV light and a multiband pass dichroic 488/561/633 for the other laser lines, which were kept for all measurements regardless of excitation lines used. These lines were individually balanced in order to obtain a similar proportion of photons from each of the dyes used in the sample; starting from the blue end of the spectra and gradually increasing the power until the expected organelle structure was seen. Emission spectra acquisition involved 28 consecutive steps using a GaAsP-PMT detector, with 10 nm bandwidth ranging 418 nm to 698 nm. During imaging, cells were maintained at 37°C under a 5 % CO_2_ atmosphere. All images were obtained such that the pixel size was in the order of half the PSF of the system (between 130nm and 100nm).

### 4.5 The Spectral Phasor Transform

The spectral phasor transform is defined as:

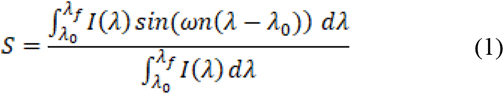

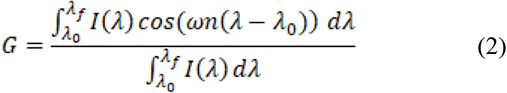

Where *I(λ)* is the photon intensity as a function of wavelength λ, collected in the spectral window [λ_0_ λ_f_]. The pulsation is chosen such that the trigonometric function’s matches the spectral window 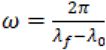. The harmonic number *n* is an integer that corresponds to the number of periods the trigonometric function are fit in the spectral window. Using equations (1) and (2), we can obtain the spectral coordinates of a spectral measurement, such that every pixel in an image maps to a position on the phasor space.

### 4.6 Higher Harmonics

One of the most important properties of the phasor transform is that it follows linear algebra vector addition [32]. This means that if in a pixel we have the spectral contribution of different components, the phasor coordinates of that pixel are the linear combination of the phasor coordinates of the components contributing to the photons in the pixel. The coefficients of the linear combinations correspond to the photon fraction of each component:

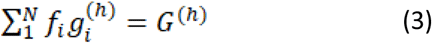

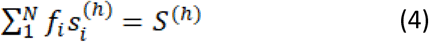

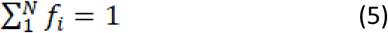

With capital (*G S*) being the experimentally measured phasor coordinates in a pixel and lower case (*g_i_ s_i_*) the phasor coordinates of the *N* components that contribute each with *f_i_* photon fraction to the measurement. The superindex *h* represents the harmonic at which the phasor coordinates are computed, it is not an exponent.

It follows that each harmonic number introduces a new independent set of equations (3) and (4). For this reason, every measurement (i.e. every pixel) has a set of 2N+1 relationships between the measured phasor coordinates and the phasor coordinates of the true components contributing to that measurement (2N+1 corresponds to the two equations (3) and (4) times the number of harmonics used N plus equation (5)). These 2N+1 equations allow one to construct a consistent system for three or fewer components with one harmonic, five or less for two harmonics, seven or less for three harmonics, and so on.

### 4.7 Pure Component Coordinates

Pure components were measured from empirical measurements with the specific dyes mentioned in the cell staining section. LAURDAN extremal phasor positions were empirically measured using the multilamellar vesicle preparations with fluid or liquid-ordered phases. For each component, cells were imaged with the respective probe and the phasor position of the probe was computed as follows: a threshold of 50 counts was applied so that only pixels with 50 or more photons were used (in the order of 10^5^ pixels survived for each component, with a total photon count in the order of 10^7^ per component). For every pixel, the phasor transform was computed at increasing harmonics, and the 2D histogram of the phasor coordinates (the phasor plot) was constructed using square bins of 0.004 phasor units per side. Based on these 2D histograms, the peak position for each distribution was located in each harmonic. See Supplementary Figure 1.

### 4.8 Phasor Unmixing

Given the phasor coordinates of the expected components, each pixel of an image is then interrogated for the fractions of each of the components. In all cases, images were phasor transformed on a pixel basis and phasor coordinates were extracted for N harmonics. In the case of solving for 4 and 5 components, N=2 harmonics were used per pixel, in the case of solving for 6 or 7 components, N=3 harmonics were used per pixel. Then the system of 2N+1 equations was constructed: equations (3) and (4) for each harmonic plus equation (5). This system of equations was solved at each pixel thus providing a set of photon fractions at each pixel.

### 4.9 Simulation Details

The emission spectra of the sample fluorophores were downloaded from Chroma spectra-viewer which provides spectral profiles at a resolution of 1nm. The chosen fluorophores were atto425, atto488, atto550, atto594, and atto647, nicely covering the typical spectral range used in fluorescence microscopy. The 4×10^5^ simulated experimental combinations of these spectra were performed on a number-of-photon basis. The number of photons in each experiment was logarithmically defined so that in the 4×10^5^ experiments, the whole range from 1 photon to 10^6^ photons was logarithmically covered, i.e. there were many more simulated experiments carried out for low photon numbers. For each experiment, a set of component fractions was generated from a uniform distribution, normalized to unity, and shuffled to prevent bias toward one component. Then, they were multiplied by the random total number of photons assigned to the specific experiment and rounded to the closest integer. This procedure produced the number of photons that were to be drawn from each component’ s spectral profile. Individual photon wavelengths were obtained in a Monte Carlo-based approach; define an area on which the pure component spectral curve is plotted, with the abscissa being the wavelength and the ordinate being the intensity as a function of wavelength. Then draw a pair of uniform random coordinates in the area and keep the abscissa if the ordinate is under the curve. This allowed us to construct a list of wavelengths which were then used to build a histogram (with a binning of 1nm) which would be the simulated experimental spectra. These spectra were then phasor transformed and unmixed with the phasor-based multiharmonic linear unmixing method, and the obtained fractions were compared to the known fractions used for each simulated experiment.

## Data and Code Availability

Raw experimental spectral images used in the figures and MATLAB scripts written for the simulations, processing of the spectral images, and figure composition are available in the public repository Figshare [33]: https://doi.org/10.6084/m9.figshare.19469828.v1

## Acknowledgements

The authors are grateful to Prof. David Jameson for proofreading the manuscript and for his valuable suggestions. LM and FI were supported in part by PEDECIBA, Agencia Nacional de Investigación e Innovación (ANII) grant FCE_3_2018_1_149047, FOCEM - Fondo para la Convergencia Estructural del Mercosur (COF 03/11) and by the Comisión Sectorial de Investigación Científica I+D (grant 369). LM is supported as Imaging Scientist by grant number 2020-225439 from the Chan Zuckerberg Initiative DAF, an advised fund of Silicon Valley Community Foundation.

