## Supplementary Material for "Phasor-Based Multi-Harmonic Unmixing for In-Vivo Hyperspectral Imaging"

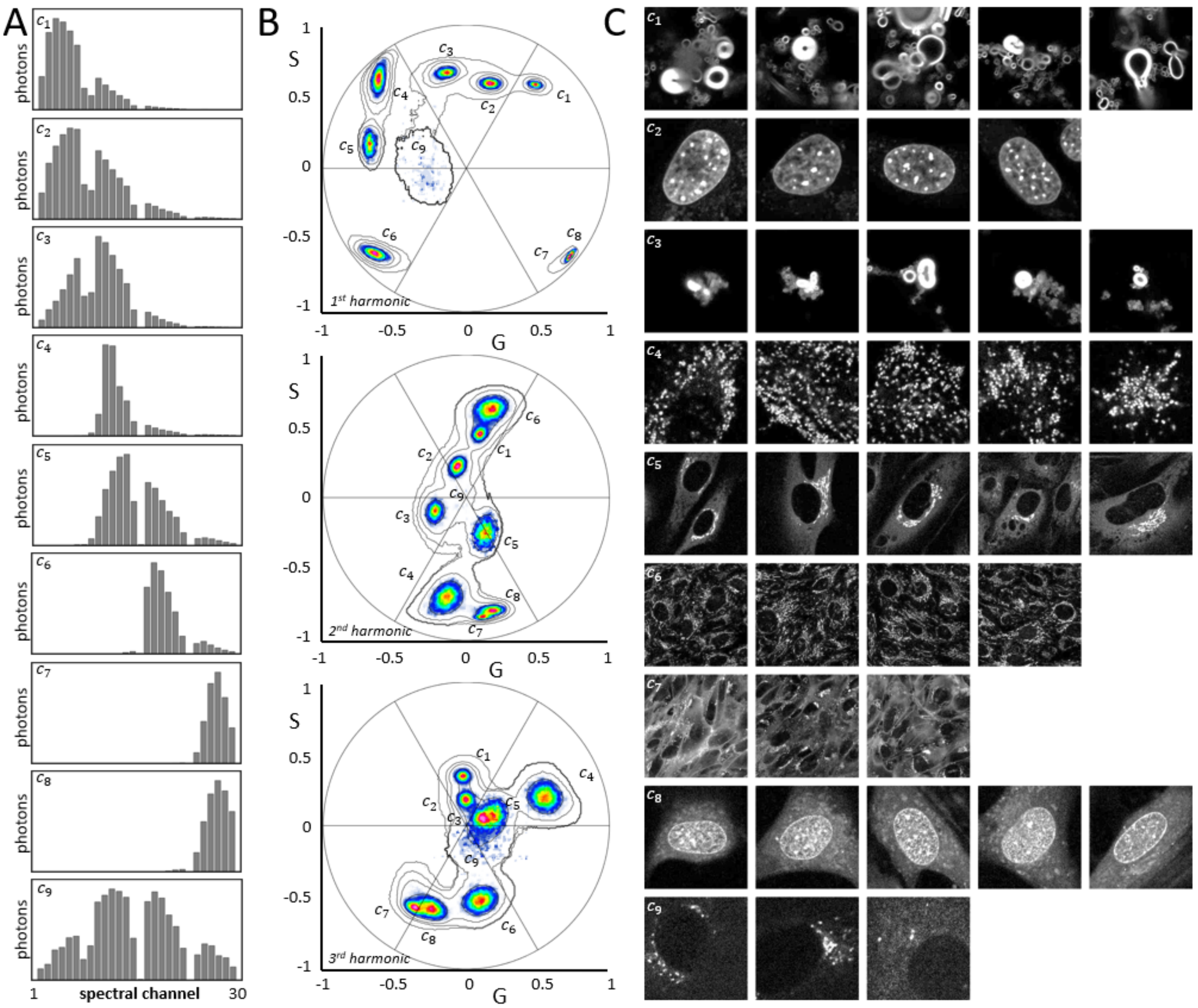


**Supplementary Figure 1. Empirical determination of the phasor positions of the pure components.** A) Accumulated spectral emission for pixels with more than 50 photon counts in each of the component image sets ($10^5$ pixels per component, $10^7$ photons per distribution). The gaps in the spectral distributions correspond to the emission dichroic band notches for the excitation laser lines. B) Spectral phasor distributions in the first three harmonics for the pixels in each set of images. C) Intensity image samples of each of the component sets from which the spectra and phasor plots where obtained. The component image sets are: ($C_1$) Non-relaxed LAURDAN, ($C_2$) Hoechst, ($C_3$) Relaxed LAURDAN, ($C_4$) LysoTracker Green, ($C_5$) Cytopainter Golgi, ($C_6$) Mitotracker Orange, ($C_7$) CellMask, ($C_8$) NucRed, and ($C_9$) unlabeled cells (autofluorescence)).

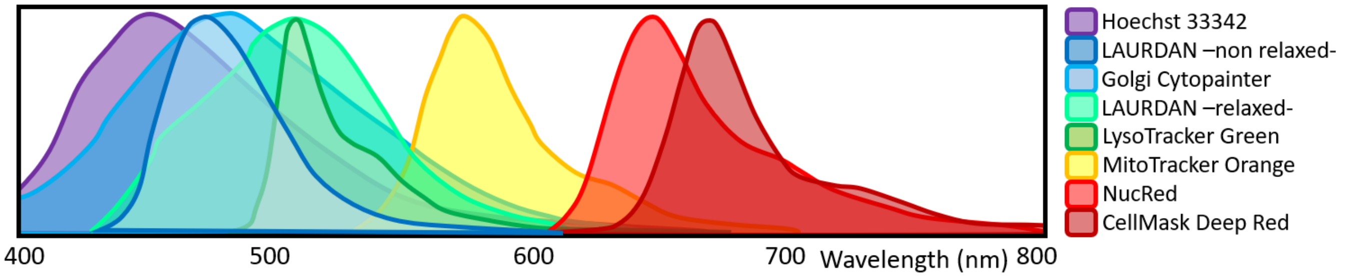


**Supplementary Figure 2. Theoretical emission spectra of the pure fluorophores.** Spectra were obtained from the respective vendors, with the exception of the two LAURDAN emissions which were obtained from [1].

[1] Malacrida L, Gratton E and Jameson D M 2015 Model-free methods to study membrane environmental probes: A comparison of the spectral phasor and generalized polarization approaches *Methods Appl. Fluoresc.* **3**

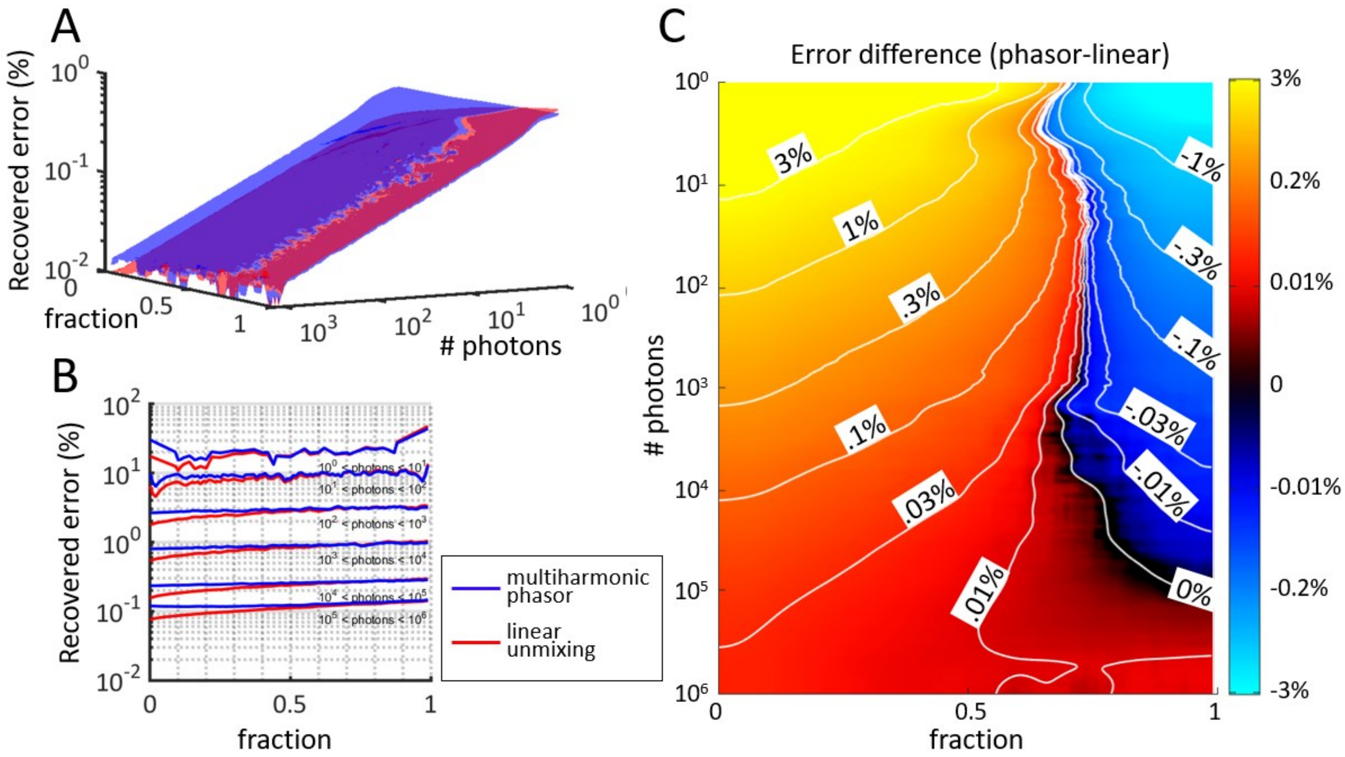


**Supplementary Figure 3. Comparison between phasor multi-harmonic unmixing and classical linear unmixing.** The two methods were compared using simulated spectra covering uniform range of fractions and logarithmic number of photons ($10^7$ simulations). A) The recovered error % (difference between ground truth and recovered fraction) is plotted for the two methods as a function of the two parameters. B) Traces of the two surfaces in (A) are plotted at different photon number regimes. C) The difference between the two surfaces shows regions where the error is greater in each method (positive for phasor error higher, negative for linear error higher).
